# Different structures and pathologies of α-synuclein fibrils derived from preclinical and postmortem patients of Parkinson’s disease

**DOI:** 10.1101/2021.11.02.467019

**Authors:** Yun Fan, Yunpeng Sun, Wenbo Yu, Youqi Tao, Wencheng Xia, Yiqi Liu, Yilin Tang, Yimin Sun, Fengtao Liu, Qin Cao, Jianjun Wu, Cong Liu, Jian Wang, Dan Li

**Affiliations:** Department of Neurology and National Research Center for Aging and Medicine & National Center for Neurological Disorders, State Key Laboratory of Medical Neurobiology, Huashan Hospital, Fudan University, Shanghai 200040, China; Interdisciplinary Research Center on Biology and Chemistry, Shanghai Institute of Organic Chemistry, Chinese Academy of Sciences, Shanghai, 201210, China; University of Chinese Academy of Sciences, Beijing 100049, China; Key Laboratory for the Genetics of Developmental and Neuropsychiatric Disorders (Ministry of Education), Bio-X Institutes, Shanghai Jiao Tong University, Shanghai, China; Bio-X-Renji Hospital Research Center, Renji Hospital, School of Medicine, Shanghai Jiao Tong University, Shanghai 200240, China; Zhangjiang Institute for Advanced Study, Shanghai Jiao Tong University, Shanghai 200240, China

## Abstract

α-Synuclein (α-syn) fibrillar aggregates are the major component of Lewy bodies and Lewy neurites presenting as the pathology hallmark of Parkinson’s disease (PD). Studies have shown that α-syn is potential to form different conformational fibrils associated with different synucleinopathies, but whether the conformation of α-syn fibrils changes in different phases of related diseases is to be explored. Here, we amplified α-syn aggregates from the cerebrospinal fluid (CSF) of preclinical (pre-PD) and late-stage postmortem PD (post-PD) patients. Our results show that compared to the CSF of pre-PD, that of post-PD is markedly stronger in seeding *in vitro* α-syn aggregation, and the amplified fibrils are more potent in inducing endogenous α-syn aggregation in neurons. Cryo-electron microscopic structures further reveal that the difference between the pre-PD- and post-PD-derived fibrils lies on a minor polymorph which in the pre-PD fibrils is morphologically straight, while in the post-PD fibrils represents a single protofilament assembled by a distinctive conformation of α-syn. Our work demonstrates structural and pathological differences between pre-PD and post-PD α-syn aggregation and suggests potential alteration of α-syn fibrils during the progression of PD clinical phases.

**Significance Statement:** Increasing evidence support different conformational α-syn fibrils in patients with different α-synucleinopathies, but whether the conformation of α-syn fibrils changes in different phases of related diseases is unknown. Here, we show that α-syn fibrils amplified from the cerebrospinal fluid (CSF) of the late-stage postmortem PD (post-PD) patient are more potent in inducing endogenous α-syn aggregation in neurons than that amplified from the preclinical (pre-PD) patient. Cryo-EM structures further reveal that the post-PD fibrils contain a novel conformation that is distinct from either the pre-PD fibrils or those previously reported. Our work suggests conformational evolution of α-syn fibrils along with PD progression.

## Introduction

Parkinson’s disease (PD), clinically presenting with akinesia, rigidity, and resting tremor, is the most common neurodegenerative movement disorder with approximately 2% of people over 65 years old affected worldwide.^1^ Lewy bodies (LBs) and Lewy neurites (LNs) that mainly comprise α-synuclein (α-syn) amyloid fibrils are the pathological hallmark of PD.^2,3^ Intracellular abnormal deposition of α-syn aggregates is also a dominant neuropathological feature of dementia with Lewy bodies (DLB) and multiple system atrophy (MSA), the other two common α-synucleinopathies.^4^ α-Syn aggregates from different α-synucleinopathies, termed as strains, demonstrate distinct pathogenic properties such as seeding and transmission features.^5-7^. Consistently, structural studies have shown that α-syn can form polymorphic fibril structures.^8-13^ These findings suggest that distinct α-syn strains may cause the clinical and pathological heterogeneities of α-synucleinopathies.

The process leading to clinically defined PD starts much earlier than that can be captured by current diagnostic criteria.^14^ Generally, the natural history of PD can be divided into three phases including the preclinical phase with no clinical signs and symptoms; the prodromal phase with the presence of non-motor symptoms and with or without slight motor symptoms; and the clinical phase that can be further classified into an early stage with cardinal motor dysfunction and a late stage with the presence of axial motor symptoms, dementia, and psychosis.^15^ Accumulating evidence supports a prion-like templating and propagation property of α-syn,^16-18^ which is believed to contribute to the distribution of α-syn pathology in patients’ brains associated with the progression of PD.^19-21^ However, given the polymorphism of α-syn fibrils, it remains unknown whether the structure of α-syn fibrils changes during spread in different phases of PD.

In the present study, we amplified α-syn aggregates in the cerebrospinal fluid (CSF) from a preclinical PD patient (pre-PD) and a late-stage postmortem PD patient (post-PD). We determined the structures of the amplified fibrils by cryo-electron microscopy (Cryo-EM) and demonstrated the structural and pathological differences between the pre-PD and post-PD fibrils. Our work prompts that the structure and pathology of α-syn fibrils may change underlying the progression of PD clinical phases.

## Results

### Clinical history of the patients

The demographic characteristics of the research subjects, including a pre-PD patient, a post-PD patient, and four healthy controls (HCs) in this study are shown in **Table 1**. CSF samples of the six subjects were collected. The CSF sample of the pre-PD patient was initially collected in 2014 as a healthy control. In 2021, as we found this patient’s CSF positive in seeding *in vitro* α-syn aggregation, we followed up this patient and clinically diagnosed him as PD. This patient has presented progressive tremor of the right hand, rigidity, and bradykinesia since 2018 (**Table 2**), indicating a preclinical phase of PD at the time of CSF sample collection. The results of non-motor symptoms (NMS) and quality of life (QoL) assessments on the pre-PD are also showed in **Table 2**. As for the post-PD patient, she was diagnosed as in the clinical late stage with severe motor symptoms and dementia on her last visit to the hospital (**Table 2**). Of note, CSF of the post-PD patient was obtained by autopsy of the postmortem patient.

**Table 1.** Demographic characteristics of the PD patients and heathy controls.

| Subjects | Sex | Age at onset (y) | Age at LP (y) | H&Y grade at LP (y) | Age at diagnosis (y) |
| --- | --- | --- | --- | --- | --- |
| Pre-PD | M | 76 | 72 | 0 | 79 |
| Post-PD | F | 50 | 70<sup>a</sup> | 5 | 50 |
| HC-1 | M | NA | 81 | NA | NA |
| HC-2 | M | NA | 56 | NA | NA |
| HC-3 | M | NA | 73 | NA | NA |
| HC-4 | M | NA | 57 | NA | NA |

LP, lumbar puncture; H&Y, Hoehn & Yahr; HC, healthy control
<sup>a</sup>Post-PD CSF was obtained by autopsy of the postmortem patient.

**Table 2.** Clinical characteristics of the PD patients in the last visit to hospital.

|  | Pre-PD | Post-PD |
| --- | --- | --- |
| Disease duration (y) | 3 | 18 |
| H&Y | 1 | 5 |
| UPDRS-III score | 12 | 57 |
| NMSS | 27 | 105 |
| GDS score | 14 | 16 |
| BDI | 16 | 23 |
| ESS | 2 | 13 |
| RBDSQ score | 2 | 7 |
| SSST-12 score | 10 | —<sup>a</sup> |
| MMSE | 30 | 12 |
| PDQ-39 | 30 | 82 |

<sup>a</sup>The post-PD patient could not accomplish the SSST-12 test
H&Y, Hoehn & Yahr; UPDRS-III, Unified Parkinson's Disease Rating Scale part III; NMSS, Non-Motor Symptoms Scale; GDS, Geriatric Depression Rating Scale; BDI, Beck Depression Inventory; ESS, Epworth Sleepiness Score; RBDSQ, Rapid-Eye-Movement Sleep Behavior Disorder Screening Questionnaire; SSST-12, Sniffin' Sticks screening 12 test; MMSE, Mini Mental State Examination; PDQ-39, 39-item Parkinson's disease questionnaire

### Pre-PD and post-PD CSF exhibit distinct abilities in seeding *in vitro* α-syn aggregation

Amplification of α-syn fibrils from the CSF of the six subjects was performed according to the protocol of the Soto lab.^22^ The thioflavin T (ThT) fluorescence was applied to monitor the aggregation kinetics of α-syn. Our result showed that the pre-PD and post-PD CSF samples, but not those of the healthy controls, were able to effectively seed α-syn aggregation *in vitro* (**Figure 1a**). Images of negative staining transmission electron microscopy (NS-TEM) confirmed the formation of amyloid fibrils in the pre-PD and post-PD samples (**Figure 1b**). Notably, the post-PD CSF was more effective than that of the pre-PD in seeding α-syn aggregation resulting in a markedly shorter lag time (**Figure 1a**). Intriguingly, the fibrils seeded with the pre-PD CSF exhibited much higher ThT fluorescence intensity than those seeded with the post-PD CSF (**Supplementary Figure 1a**). Consistently, more fibrils in the pre-PD CSF-seeded sample were observed by NS-TEM (**Figure 1b**). Furthermore, we measured the concentration of soluble α-syn remained in the supernatant of each sample, and calculated the conversion rate of α-syn monomer to fibrils. In line with the ThT and NS-TEM data, significantly more α-syn molecules converted to fibrils when seeded with the pre-PD CSF compared to that seeded with the post-PD CSF (**Supplementary Figure 1b**). Taken together, our data demonstrate distinct abilities of the pre-PD and the post-PD CSF in seeding α-syn aggregation: the former results in more aggregation, while the latter results in faster aggregation.

**Figure 1.**
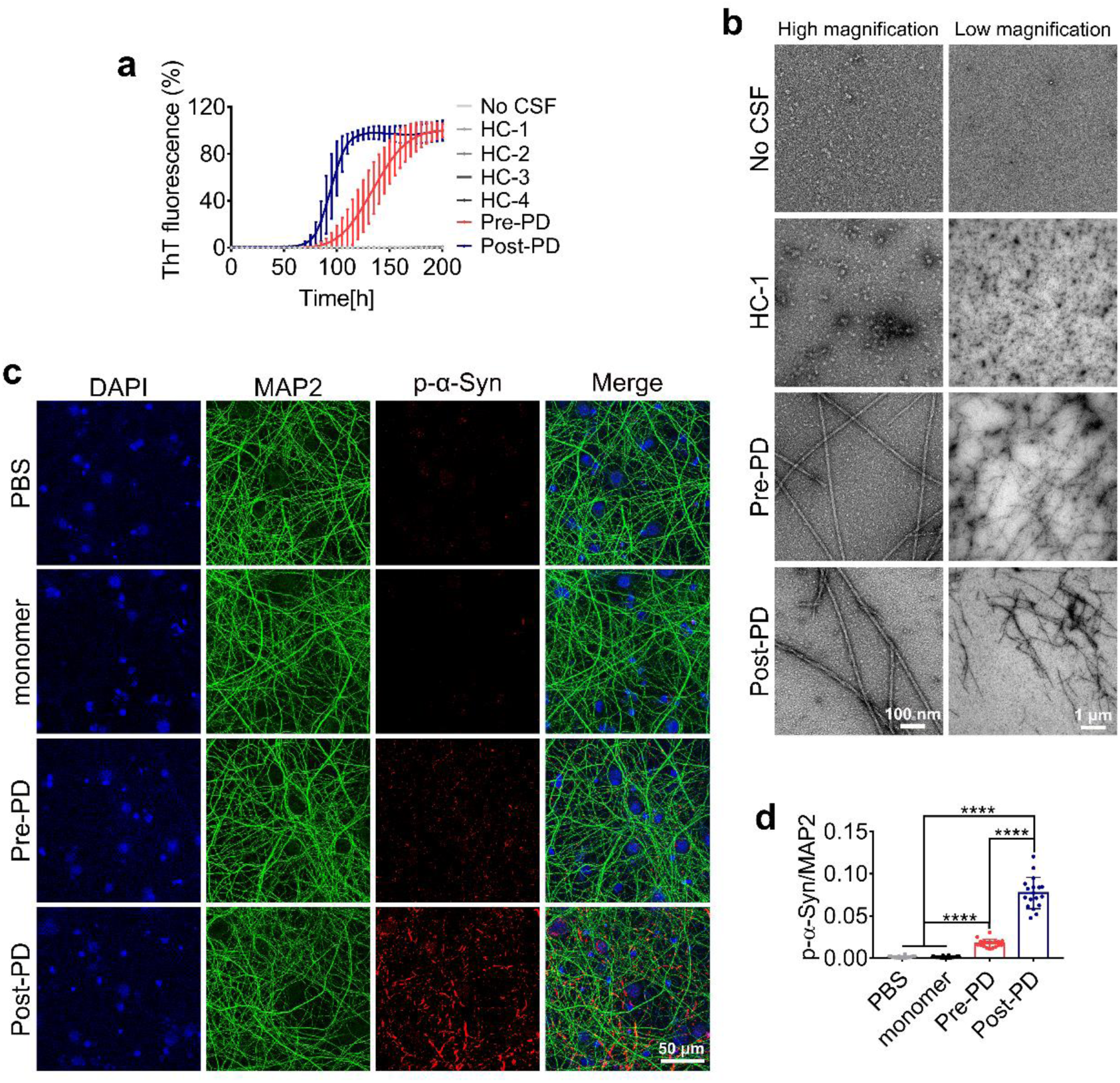
Different pathological properties of the CSF and CSF-amplified α-syn fibrils from pre-PD and post-PD patients. **a**. ThT kinetic assay for α-syn aggregation seeded with CSF from healthy controls (HCs), a pre-PD patient and a late-stage post-PD patient. Each sample contains 40 μl CSF and 160 μl reaction solution of α-syn monomer (50 μM). ThT intensity was expressed as a percentage of the maximum fluorescence intensity. Data shown are mean ± SD, n=3 individual experimental samples. **b**. NS-TEM imaging of the samples at the end of the ThT assay in (**a**). Images were taken at high and low magnification, respectively. **c**. Confocal microscopic images of rat cortical primary neurons treated with PD-derived α-syn PFFs. Primary neurons at 7 days were treated with 100 nM pre-PD or post-PD PFFs for 22 days. The fixed neurons were immunostained for DAPI (blue), S129-phosphorylated α-syn (p-α-syn, red) and MAP2 (green). **d**. Quantification of p-α-syn in (**c**) as expressed by p-α-syn intensity normalized to MAP2 intensity. Data are shown as mean ± SD of 18 images in three independent experiments. Statistical significance was measured by one-way ANOVA followed by Tukey’s post-hoc test. \*\*\*\**P* < 0.0001.

### α-Syn fibrils amplified from the post-PD CSF are less stable and more pathological to primary neurons

To characterize whether the fibrils amplified from pre-PD and post-PD CSF respectively are similar or not, we first treated the two CSF-derived α-syn fibrils with different concentrations of proteinase K (PK) at 37 °C for half an hour. The result revealed that the post-PD fibrils were digested faster than the pre-PD fibrils (**Supplementary Figure 1c**), indicating that the post-PD fibrils are less stable than the pre-PD fibrils.

Next, we sought to investigate the pathological property of the fibrils to primary neurons. Firstly, we equalized the fibril amounts in each sample and sonicated the fibrils to prepare preformed α-syn fibril seeds (PFFs) (**Supplementary Figure 1d**). The PFFs in each sample were of similar lengths determined by atomic force microscopy (AFM) (**Supplementary Figure 1e, f**). An equal amount of PFFs in each sample was confirmed by SDS-PAGE before treating neurons (**Supplementary Figure 1g**). Next, we treated rat cortical primary neurons with the PFFs. The result showed that in comparison with phosphate buffered saline (PBS) or α-syn monomer, both pre-PD- and post-PD-derived PFFs can efficiently seed endogenous α-syn aggregation (**Figure 1c, d**). Moreover, the post-PD PFFs were significantly more potent in inducing α-syn pathology in primary neurons than the pre-PD PFFs (**Figure 1c, d**). Collectively, these results suggest that the α-syn fibrils amplified from the post-PD CSF feature lower stability and higher pathology than those from the pre-PD CSF.

### Cryo-EM structures of α-syn fibrils amplified from the pre-PD CSF

To investigate the structural basis underlying the distinct properties between the pre-PD- and post-PD-derived fibrils, we performed cryo-EM to determine their atomic-level structures. The condition of cryo grids and the concentration of fibrils were optimized to acquire qualified high-resolution cryo-EM micrographs for structure determination (**Figure 2a**). Subsequently, we conducted a series of processing routines for the micrographs by the RELION software^23^ (**Table 3**). Micrograph processing after 2D classification revealed that the pre-PD fibrils consist of two polymorphs, termed as Type 1A and Type 2, occupying approximately 85% and 15% of the picked fibrils, respectively (**Figure 2a, b**). After helical reconstruction in RELION, we obtained the 3D density map of Type 1A fibrils with an overall resolution of 2.78 Å (**Figure 2c ,Supplementary Figure 2**). Type 1A fibril presents as a left-handed helix with a width of ∼12 nm and a half helical pitch of ∼78 nm (**Figure 2d**). The two protofilaments in Type 1A fibril intertwine along an approximate two-fold screw axis. The helical twist between neighboring α-syn subunits is 179.45° with a helical rise of 2.44 Å (**Table 3**). The Type 2 fibril also comprises two protofilaments; however, it appears to be straight, which prevents the 3D reconstruction by the current technology of cryo-EM (**Figure 2a, b**).

**Figure 2.**
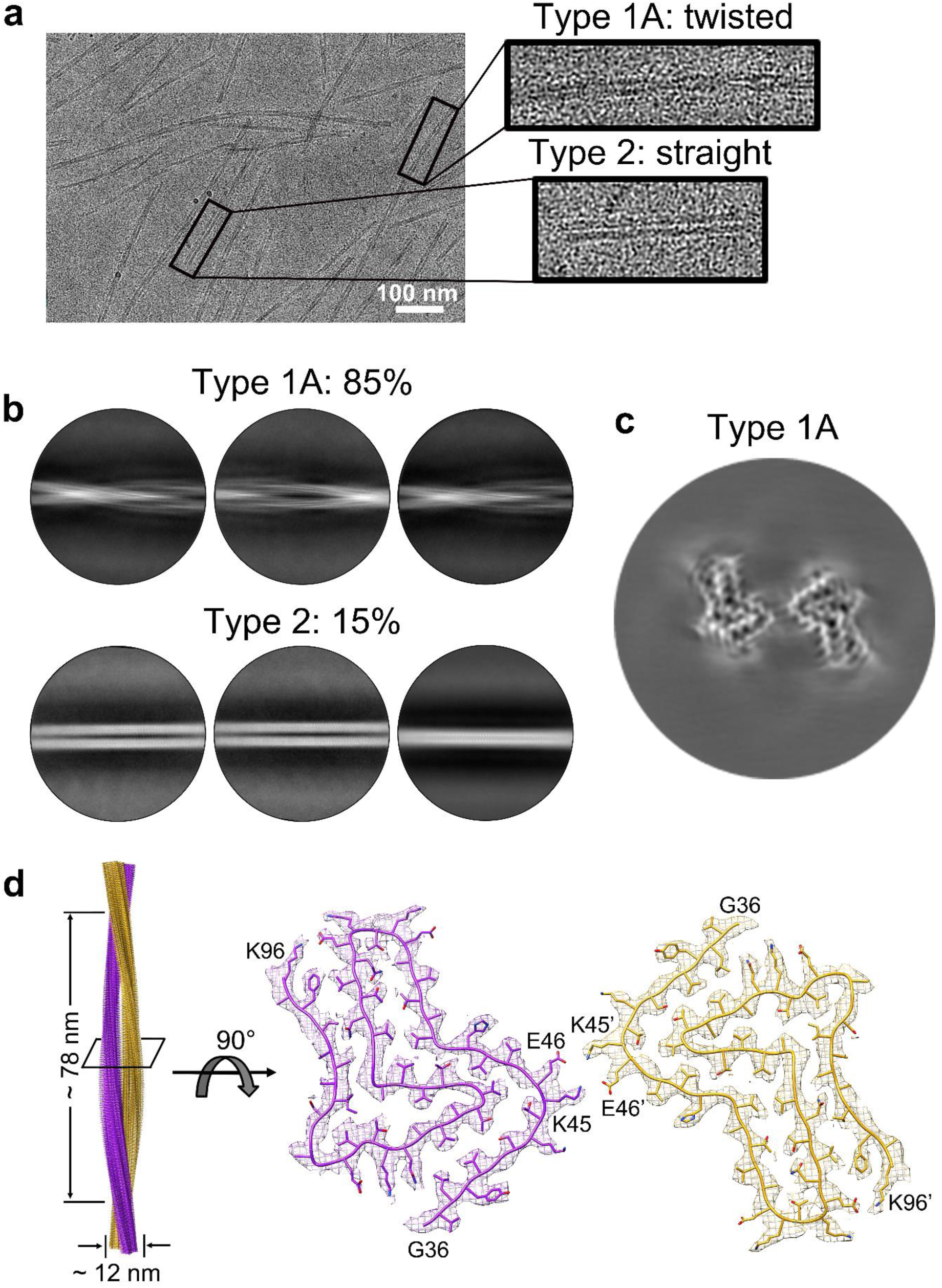
Cryo-EM structure determination of α-syn fibrils amplified from the pre-PD CSF. **a**. Cryo-EM micrographs of the pre-PD CSF amplified fibrils. Zoom-in images show the Type 1A and Type 2 polymorphs. Scale bar =100 nm. **b**. 2D class average of Type 1A (top) and Type 2 (bottom) fibrils in the box size of 686 pixels. The percentage of Type 1A and Type 2 fibrils in the picked fibrils is indicated. **c**. 3D density map of the Type 1A fibril. **d**. Cryo-EM reconstruction density map of the Type 1A fibril with fibril width and half pitch labeled (left). Cross-section view of the density map with the atomic model fitted in it (right). The two protofilaments in the Type 1A fibril are colored in purple and yellow, respectively. The figure was prepared by using UCSF Chimera v1.13.

**Table 3.**
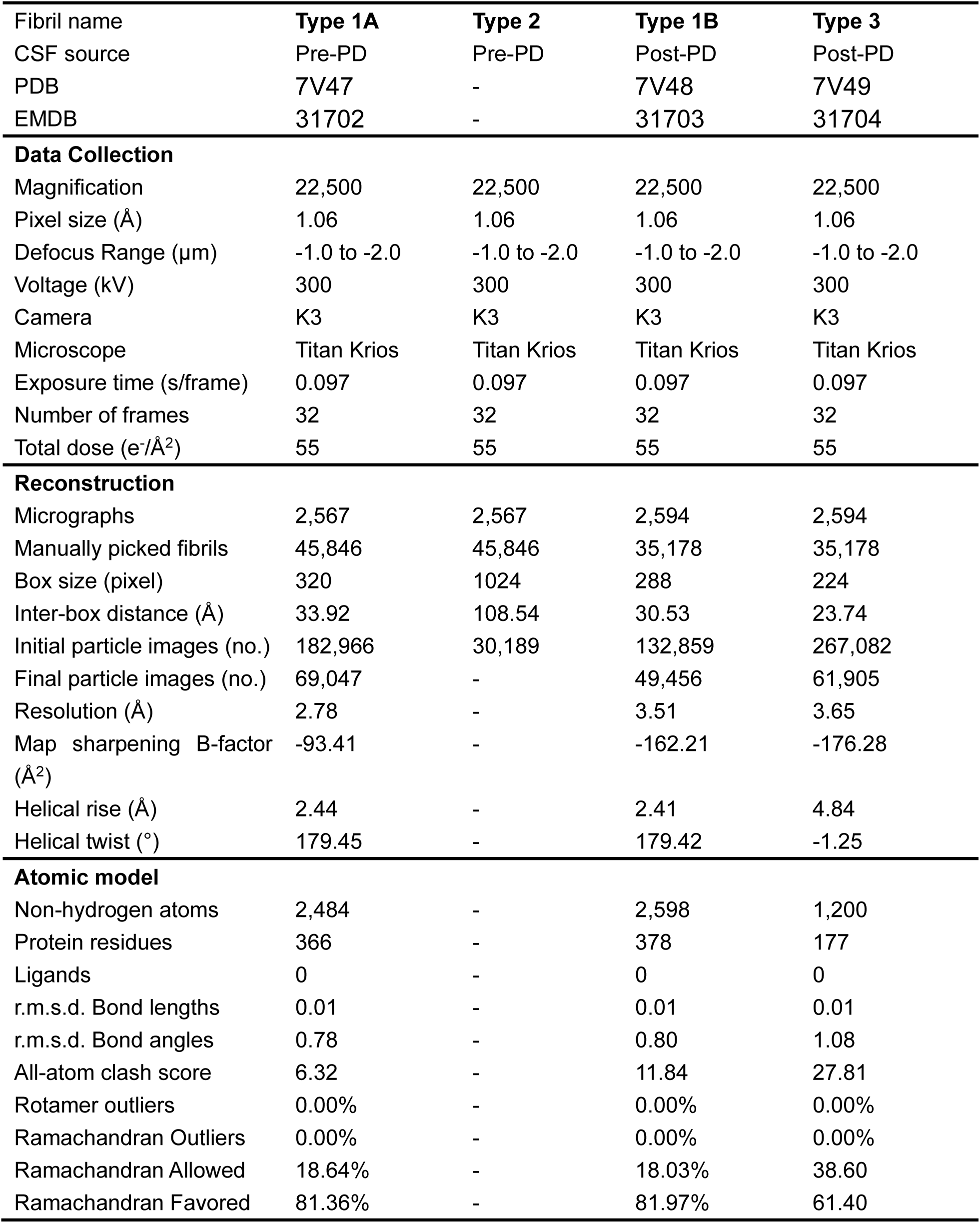
Cryo-EM structure determination and model statistics.

Based on the density map of Type 1A, we were able to unambiguously build an atomic structure model for the Type 1A fibril comprising residues 36-96 (**Figure 2d**). Two pairs of salt bridges formed by K45 and E46 hold the two protofilaments of Type 1A fibrils together. The structure of α-syn monomer in the Type 1A fibril is similar to that in the so-called polymorph 2b fibril (PDB ID: 6SST), an *in vitro* assembled α-syn fibril, with an RMSD of 3.727 Å over 61 C-α atoms^10^ (**Supplementary Figure 3**). A major difference between Type 1A and polymorph 2b is that residues 14-25 in polymorph 2b are ordered and form a steric-zipper-like interactions with residues 85-91, while this additional N-terminal region is flexible and not able to be determined in Type 1A (**Supplementary Figure 3**).

### Cryo-EM structures of α-syn fibrils amplified from the post-PD CSF

Similar to the fibrils amplified from the pre-PD CSF, those from the post-PD CSF also consist of two polymorphs, termed as Type 1B and Type 3, occupying approximately 72% and 28% of the total picked fibrils, respectively (**Figure 3a ,b**). After helical reconstruction in RELION, we obtained 3D density maps for Type 1B and Type 3 with an overall resolution of 3.51 Å and 3.65 Å, respectively (**Figure 3c ,Supplementary Figure 2**). The Type 1B fibril presents as a left-handed helix with a width of ∼11 nm and a half helical pitch of ∼75 nm (**Figure 3d**). It consists of two protofilaments intertwining along an approximate two-fold screw axis. The helical rise between neighboring α-syn subunits is 2.41 Å and the helical twist is 179.42° (**Table 3**). The overall structure of α-syn in the Type 1B fibril is nearly identical to that in the Type 1A fibril (**Supplementary Figure 3**).

**Figure 3.**
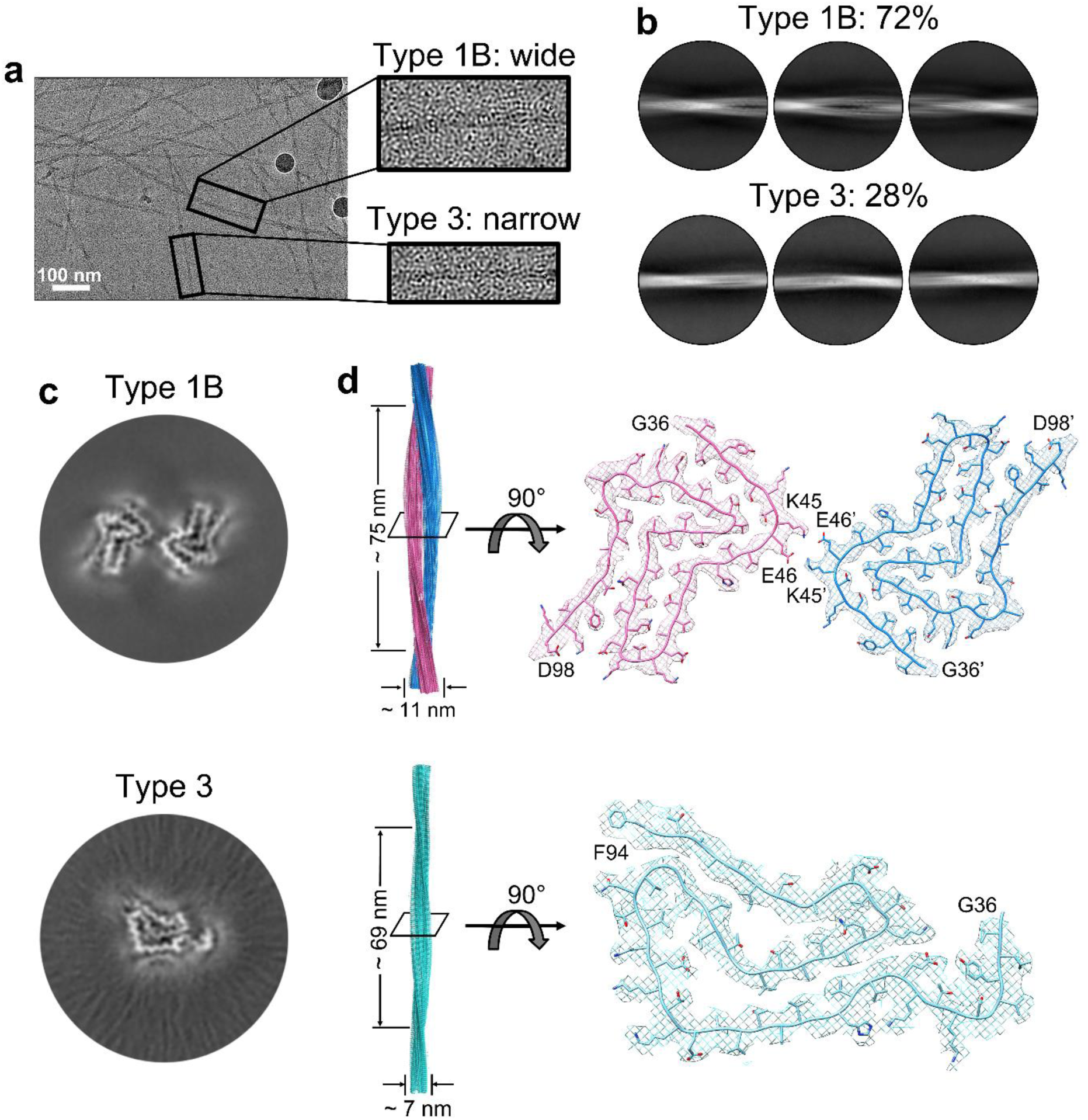
Cryo-EM structure determination of α-syn fibrils amplified from the post-PD CSF. **a**. Cryo-EM micrographs of the post-PD amplified fibrils. Zoom-in images show the Type 1B and Type 3 polymorphs. Scale bar =100 nm. **b**. 2D class average of Type 1B (top) and Type 3 (bottom) fibrils in the box size of 686 pixels. The percentage of Type 1B and Type 3 fibrils is indicated. **c**. Cryo-EM 3D density maps of the Type 1A (top) and Type 3 (bottom) fibrils. **d**. Cryo-EM reconstruction density maps of the Type 1B (left, top) and Type 3 (left, bottom) fibrils with fibril width and half pitch labeled. Cross-section view of the density maps with the atomic model fitted in it (right). The two protofilaments of the Type 1B fibril are colored in pink and blue, respectively. The figures were prepared by using UCSF Chimera v1.13.

Distinctively, the Type 3 fibril contains only one protofilament (**Figure 3b ,c, d**). The 3D reconstruction revealed that Type 3 features a half pitch of ∼69 nm and a fibril width of ∼7 nm (**Figure 3d**). The twist angle and helical rise between neighboring subunits is -1.26° and 4.84 Å, respectively (**Table 3**). According to the density map, we built a structure model for the Type 3 fibril which is composed of residues 36-94 forming eight β sheets (**Figure 3d ,Supplementary Figure 4**). The structure of the Type 3 α-syn fibril is different from any α-syn fibril structures reported so far, representing a new polymorph of α-syn fibrils.

### Structural comparisons highlight the pathological relevance of the Type 3 fibril

Given the high similarity between the Type 1A and Type 1B fibril structures, the difference between the biochemical and pathological properties of the pre-PD- and post-PD-derived fibrils is most likely contributed by the Type 3 fibril. Although Type 3 represents as a distinctive polymorph of α-syn fibrils, the structure of α-syn molecule in the fibril presents similarity with that in some known α-syn fibril polymorphs. For example, the structures of α-syn molecules in Type 3 and the so-called polymorph 1a (*in vitro* assembled, PDB ID: 6A6B) fibrils show obvious similar folding topologies, although their structure similarity is low with an RMSD of 4.85 Å over 58 C-α atoms (**Supplementary Figure 5a)**. Similar folding of α-syn also presents in the fibrils derived from MSA patients. Cryo-EM structures of α-syn fibrils directly purified from the brains of MSA patients showed two types (type I and type II) of fibril polymorphs in the *ex vivo* fibrils.^12^ Although the structures of α-syn molecules in each polymorphs as well as in each protofilament of the same polymorph are different, they share a conserved structural motif formed by residues K43-F94/Q99 (**Supplementary Figure 5b**). Except for the conserved structural motif, the rest of α-syn structures in the MSA polymorphs are heterogeneous and seem to form due to the presence of co-factors and posttranslational modifications (PTMs) (**Supplementary Figure 5b**).^24^ Indeed, in the absence of native co-factors and PTMs, the *in vitro* assembled α-syn fibrils seeded by the *ex vivo* fibrils of MSA present a polymorph, in which α-syn folds into the nearly identical structure as the conserved structural motif in the *ex vivo* fibrils (**Figure 4a**).

**Figure 4.**
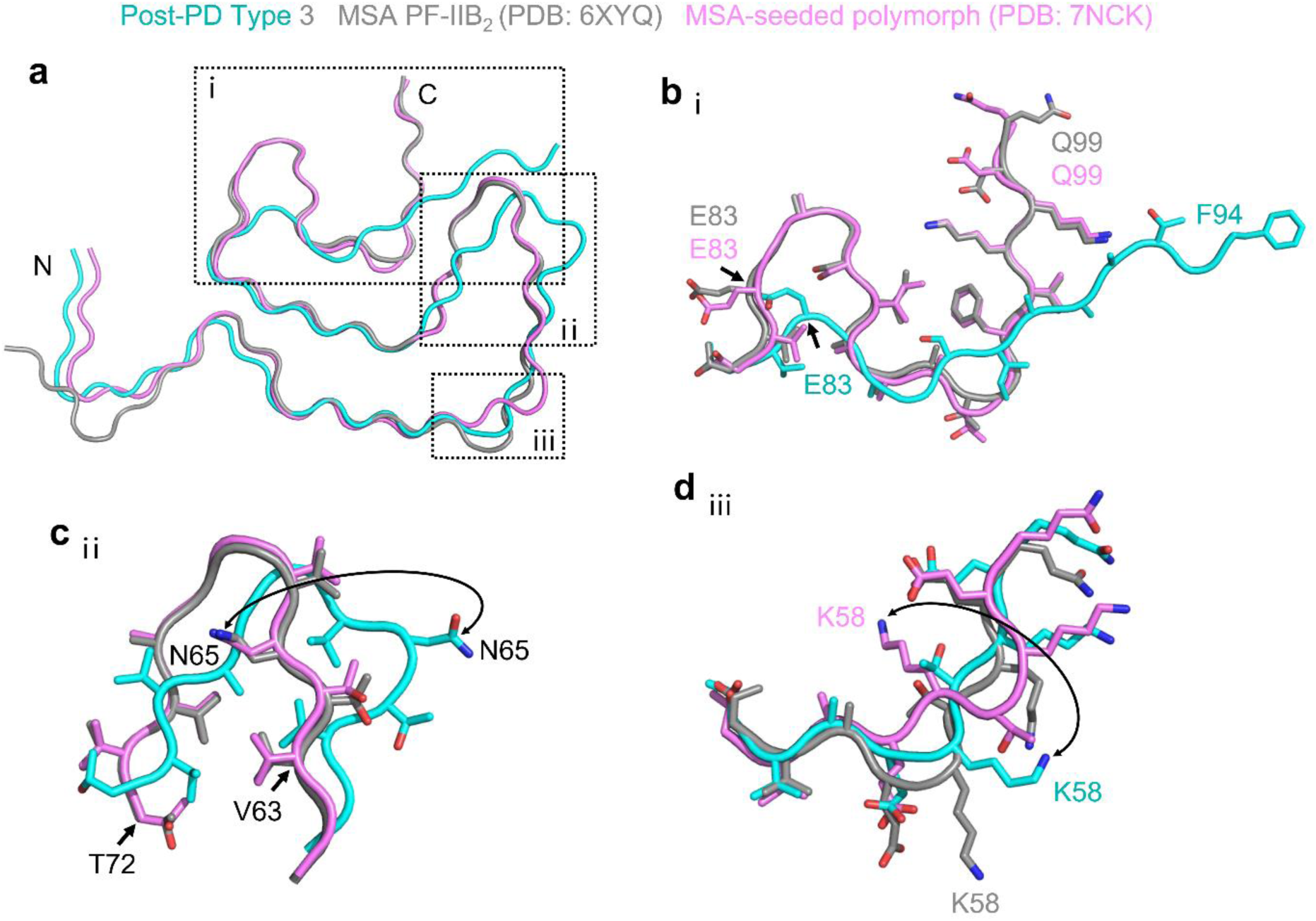
Structural comparison of the Type 3 fibril with MSA-derived polymorphs. **a**. Alignment of α-syn structures in the Type 3 fibril (cyan), the conserved structural motif in MSA-purified fibrils (grey), and the *in vitro* formed fibril seeded with MSA-purified fibrils (purple). The major differences of the structures are shown in the zoom-in views in (**b-d**). Structural differences of C-terminal (**b**) and residues 59–72 (**c**) between Type 3 fibrils and MSA-purified fibrils or MSA *in vitro* seeded fibrils are shown. **d**. Directions of K58 side chains in the three structures. Side chains in **b**-**d** are shown as sticks.

Intriguingly, α-syn in the Type 3 fibril amplified from the post-PD CSF exhibits a similar fold to that in the MSA-derived fibrils (**Figure 4a**), although their structure similarities are low with an overall RMSD over 59 C-α atoms (residues 36-94) ranging from 5.047 to 8.147 Å. One of the structural differences lies in the C-terminal region. Instead of folding into a short L-shaped motif in the MSA-derived fibrils, the C-terminal region of the Type 3 fibril (residues 83-94) appears more extended (**Figure 4b**). Another structural difference lies in residues 59–72, which form a conserved turn in the MSA-derived fibrils, while form a β hairpin in the post-PD-derived Type 3 fibril with the side chain of residues 65-72 flipped to the other side (**Figure 4c**). In addition, the side chain of K58 in the Type 3 fibril pointing toward the solvent, which is distinct from that in the MSA-seeded fibril, but similar to that in the MSA *ex vivo* fibrils (**Figure 4d)**. In the MSA *ex vivo* fibrils, K58 points outwards either interacting with co-factors or the opposite α-syn subunit^12,24^; similarly, in the Type 3 fibril, K58 might also interact with residual unknown co-factors in the CSF since CSF represents 20% v/v of the fibril sample. Collectively, these fibril structures support a pathological significance of polymorph Type 3 and indicate that the structure of α-syn in the Type 3 fibril might partially present in the PD patient.

## Discussion

The clinical manifestations of PD present a great variability along the disease progresssion.^15^ Closely associated with the clinical phenotypes and disease progression of PD, it has been suggested that the abnormal deposition of α-syn aggregates spreads subsequently from the olfactory bulb and medulla oblongata, then to the midbrain and basal forebrain, and finally to the neocortex.^19,20^ However, it remains elusive whether the conformation of α-syn aggregates changes along its propagation in the advance of disease. Here, we investigated the CSF collected from two PD patients in different disease phases: one is at the preclinical stage and the other is at the end stage. We found that the pre-PD and post-PD CSF behaved differently in seeding *in vitro* formation of α-syn fibrils. We further investigated the ability of the amplified fibrils to induce α-syn pathology in rat primary neurons and determined their cryo-EM structures. Our results demonstrate that the post-PD-derived α-syn fibrils are more pathological than the pre-PD-derived fibrils, and the post-PD fibrils contain a distinctive fibril polymorph, in which α-syn folds into a similar topology to that in the MSA-derived fibrils.

Note that the fibrils amplified from PD CSF are least possible to fully represent those in the CSF or brain of PD patients. The fibrils directly purified from patient brains usually contains co-factors and PTMs.^12,25-27^ The *in vitro* formed fibrils in the absence of native modifications thus can hardly fully recapitulate the structures of *in vivo* fibrils. However, the presence of patient’s biological fluid and tissue samples such as CSF and brain homogenate may seed the *in vitro* formation of disease-related fibrils. How much the seeded fibrils would represent their mother fibrils is partially dependent on the conditions. In our work, we adopted a so-called protein misfolding cyclic amplification (PMCA) protocol established by the Soto lab for the α-syn fibril amplification from PD CSF.^22^ Our results showed that the CSF of age-matched healthy controls yielded no fibrils; in contrast, the CSF of PD patients, both pre-PD and post-PD, efficiently stimulated fibril formation. This indicates that the obtained α-syn fibrils are closely related to the fibril seeds in the PD CSF. Meanwhile, we notice two structural studies of α-syn fibrils amplified from PD brain homogenates post on *BioRxiv* recently.^13,28^ Interestingly, in one study, the reported α-syn fibril structures are very similar to our Type 1a and 1b fibrils.^28^ While, in the other study that applied four short cycles rather than one long cycle of amplification, the reported fibril structures are distinctive from any known wild-type α-syn fibril structures but similar to that formed by Y39-phosphorylated α-syn^13,29^, which may reflect the existence of pY39-like fibril conformation in the PD brain. Taken together, the amplified fibrils from patients biological fluid and tissue samples may partially represent their mother fibrils. The *in vitro* fibril amplification cycles and conditions may influence which fibril conformation is amplified.

In this work, we used two PD patients in different clinical phases, and observed structural and pathological differences between their CSF. These differences may result from the conformational change of α-syn fibrils during disease progression, but may also simply reflect individual differences. However, based on the current knowledge, individual difference is less likely to result in such a large difference in fibril conformations. The extensive structural studies of Tau fibrils purified from postmortem brains have shown that different patients of the same tauopathy have very similar polymorphs of Tau fibrils.^25-27,30-32^ In contrast, patients from different tauopathies have distinct polymorphs of Tau fibrils.^25-27,31,33^ Consistent with Tau studies, different patients of MSA also present similar α-syn fibril polymorphs.^12^ These studies suggest that individual difference is marginal to the development of fibril polymorphs. Thus, the structural difference between pre-PD and post-PD derived fibrils demonstrated in this work is plausible to reflect the conformational change of α-syn fibrils during disease progression.

## Materials and Methods

### Clinical assessments

This study was approved by the Human Studies Institutional Review Board, Huashan Hospital, Fudan University. All subjects provided their written informed consent to clinical assessments and lumber puncture for CSF collection. The post-PD patient died two years after the last following-up and her postmortem CSF samples were obtained from the Body Donation Station in Fudan University (Shanghai Red Cross Society). All procedures conducted in the study were in conformity with the ethical standards of Declaration of Helsinki.

Four healthy controls (HCs) were free of any known neurodegenerative disease both at their baseline visit and the last following-up. The two PD patients were diagnosed by two neurologists that specialize in movement disorders. The diagnosis was according to the Movement Disorder Society Clinical Diagnostic Criteria^14^ and underwent a series of clinical evaluations comprising of demographic information, motor and non-motor symptoms, and quality of life (QoL). The Unified Parkinson’s Disease Rating Scale-part III (UPDRS-III) and the Hoehn and Yahr (H&Y) scale were used to assess motor symptoms. Non-motor symptoms were evaluated by the Non-Motor Symptom Questionnaire (NMSQ).^34^ Both Geriatric Depression Scale (GDS) and Beck Depression Inventory (BDI) were applied to assess the depression.^35,36^ Two sleepiness problem, rapid-eye movement (REM)-sleep behavior disorder (RBD) and excessive daytime sleepiness (EDS) were evaluated by the REM-sleep Behavior Disorder Screening Questionnaire (RBDSQ)^37^ and the Epworth Sleepiness Scale (ESS)^38^, respectively. The sniffin’ sticks screening 12 test (SSST-12) was used to assess the olfaction function. The Mini-Mental State Examination (MMSE) was applied to evaluate global cognitive function.^39^ QoL was measured by the 39-item Parkinson’s Disease Questionnaire (PDQ-39).^40^

### CSF collection and handing

Except for the post-PD whose CSF was obtained in autopsy, CSF samples of other subjects were collected by lumbar puncture as we previously described^41^. In brief, after fasting of no less than 8 h, lumbar puncture was performed at the interspace of L4/L5 with an atraumatic needle after local anesthesia. First approximately 2 ml CSF was for routine CSF assessment, and last 8-10 ml CSF was collected in polypropylene tube for biobanking. All intra-vitam CSF samples were centrifuged at 2,000 g for 10 min at 4 °C. The postmortem CSF sample of the post-PD was centrifuged at 20,000 g for 20 min at 4 °C within 1h. After centrifugation, the supernatants were aliquoted and immediately frozen in liquid nitrogen, followed by storing at -80 °C until use.

### Expression and purification of recombinant α-syn monomer

The preparation of α-syn monomer was performed as we previously reported with minor modifications.^8^ In brief, gene expressing human wild type full length α-syn was ligated into the pET22 vector. The recombinant plasmid was transformed into E. coli BL21(DE3) cells with the co-transformation of fission yeast N-acetyltransferase complex B to express N-terminally acetylated α-syn.^42^ E. coli BL21(DE3) cells were treated with 1 mM isopropyl-1-thio-D-galactopyranoside (IPTG) at 25 °C for 10 h to induce the expression of α-syn. The bacterial pellets were collected and lysed in the buffer containing 100 mM Tris-HCl (pH 8.0), 1 mM EDTA, 1 mM phenylmethylsulfonyl fluoride. After crushed through the high-pressure cell crusher, the solution was centrifuged at 14000g at 4 °C for 30 min. The supernatant was boiled for 10 min followed by centrifugation. To remove nucleic acids, 10 mg/ml streptomycin was added to the supernatant with continuous stirring at 4 °C for 30 min. After centrifugation, the supernatant was treated with 2-6 M HCl to adjust the pH value to 3.5, followed by centrifugation again. Then the supernatant was dialyzed in 25 mM Tris-HCl (pH 8.0) at 4 °C overnight. After dialysis, the sample was filtered through a 0.22 μm filter and then purified by the Q column (GE Healthcare, 17-5156-01) and the Superdex 75 column (GE Healthcare, 28-9893-33). The concentration of the purified α-syn was detected by the bicinchoninic acid (BCA) assay kit (Pierce). The purity of the α-syn was determined by SDS–PAGE.

### ThT kinetic assay

The amplification of α-syn aggregates in CSF were performed according to the protocol developed by Soto Lab.^22^ To remove any α-syn aggregates potentially formed in the process of freeze-thaw, purified α-syn monomer was thawed and then filtered through a 100 kDa cutoff filter (PALL, OD100C35) by centrifuged at 9000g for 10 min at 4 °C. For CSF seed reactions, 40 μl CSF and 160 μl of the reaction mix to give final concentrations of 100 mM piperazine-N,N’-bis (ethanesulfonic acid) (PIPES, pH 6.5), 500 mM NaCl, 50 μM α-syn monomer, 10 μM ThT, and 0.05% NaN3 were added to the wells of a black 96-well plate with a clear bottom (ThermoFisher, 165305). The No CSF sample was performed under the same conditions. Samples were incubated at 37 °C in the BMG FLUOstar Omega plate reader with cyclic agitation of 1min shaking (500 rpm, orbital) followed by 29 min rest. The amplification kinetics curve of the CSF seeded α-syn assembly reaction system, represented by ThT fluorescence, was measured at excitation of 440-10 nm and emission of 480-10 nm every 30 min.

### Negative staining transmission electron microscopy (NS-TEM)

5 mL of sample solution was loaded onto the glow-discharged 200 mesh carbon coated copper grids (Beijing Zhongjingkeyi Technology Co., Ltd.). After incubations of 45 s, the sample was removed by the filter paper, and the grid was washed with 5 μl double-distilled water and 5 μl 2% w/v uranyl acetate sequentially. Then another 5 μl 2% w/v uranyl acetate was applied to stain the sample for 45 s. After removing the excess buffer, the infrared lamp was used to further dry the grid. The sample imaging was accomplished by a Tecnai T12 microscope (FEI Company) operated at 120 kV.

### Atomic force microscopy (AFM)

The AFM was applied to determine the size distribution of α-syn PFFs. All samples were diluted to 10 μM with the buffer containing 100 mM PIPES (pH 6.5), 500 mM NaCl. 5 μL of the diluted sample was added to the freshly cleaved mica surface and incubated for 5 min at room temperature (RT). Unbound PFFs were then softly washed away by double-distilled water, followed by using nitrogen flow to dry the sample. Images were collected by the Nanoscope V Multimode 8 (Bruker) in the ScanAsyst air mode. Sample scanning was performed using the SCANASYST-AIR probe with force constant of 0.4 N/m and resonance frequency of 70 kHz (Bruker). Images were obtained at 512 × 512 pixels with scan rate at 1.44 Hz. The size distribution of the PFFs were analysed using the Nanoscope Analysis software (version 1.5, Bruker).

### Concentration measurement of α-syn fibrils/PFFs

Fibril samples were centrifuged at 14,462 g for 1 h at 25 °C. The resultant supernatants were carefully removed to another 1.5 ml microtube. The protein concentration in the supernatant was measured by the BCA assay (Pierce) according to the manufacturer’s instruction. The quantity of protein fibrils in the pellet was calculated as the quantity of total monomer minus the quantity of soluble protein in the supernatant. Then the pellets were washed twice and resuspended in PBS to the indicated concentration for the following experiments.

To prepare PFFs, equal concentrations of resuspended fibrils from different samples were sonicated for 15 cycles (1 s on/off per cycle, 20% amplitude, JY92-IIN sonicator) on ice. The equal concentration of the PFFs in different samples were confirmed by SDS–PAGE stained by Coomassie brilliant blue. In brief, PFFs were diluted in the buffer containing 50 mM Tris (pH 8.0),150 mM NaCl, 1% Triton X-100, 2% SDS, followed by sonication for 5 min and boiling for 30 min. Add SDS-loading buffer to the sample and boiled for another 10 min to fully dissolve the PFFs. 4%-20% Bis-Tris gels were used. Intensities of the protein bands on the gel were analyzed using Image Lab 3.0 (Bio-Rad).

### Proteinase K (PK) digestion assay

PK digestion was carried out as previous reported.^43^ Briefly, a gradient concentration (0.125 to 2 μg/ml) of PK was added to 5 μg of α-syn PFFs. The reaction was incubated at 37 °C for 30 min. The digestion reaction was stopped by heating the samples up to 100 °C for 10 min to inactivate PK. The samples were mixed with the same volume of 2X SDS-loading buffer and then boiled for 10 min. The digested products were determined by western blotting using anti-α-syn clone 42 (BD Biosciences).^44^

### Rat primary cortical neuronal culture

Pregnant Sprague-Dawley (SD) rats were commercially obtained from Shanghai SIPPR BK Laboratory Animals Ltd, China. Rat primary cortical neurons were prepared from the manually separated cortex of embryonic day (E) 16–E18 SD rat embryos according to a previous described protocols^8^. Briefly, primary neuron suspensions comprising 1.5 × 10^5^ cells were added to the 24-well plate that pre-covered with poly-D-lysine coated coverslips. Primary neurons were cultured in Neurobasal media supplemented with B-27 supplement, 0.5 mM L-glutamine and 1% penicillin-streptomycin. Primary neurons cultured at 7 days (DIV7) were treated with PFF samples for 22 days and then collected and fixed for immunostaining.

### Immunofluorescence imaging

After removing the Neurobasal media, primary neurons cultured on coverslips at DIV29 were washed with PBS containing 1 mM CaCl2 and 5 mM MgCl2 and then fixed using fixation solution consisting of PBS, 8% paraformaldehyde and 4% sucrose for 10 min at RT. Neurons were treated with PBS containing 0.15% Triton X-100 for permeabilization, followed by blocking with 5% BSA dissolved in PBS for 30 min at RT. After that, primary neurons were incubated with anti-p-α-syn (Abcam, ab51253,1:1,000 dilution) and anti-MAP2 (Abcam, ab5392, 1:1,000 dilution) antibodies at 4 °C overnight. After washing three times using PBS, secondary antibodies including Alexa Fluor 488-conjugated goat anti-chicken and Alexa Fluor 568-conjugated goat anti-rabbit secondary antibodies (Life Technologies, A11039 and A11036, respectively, 1:1,000 dilution) were incubated with primary neurons for 1 h at RT. After mounting with ProLong Gold Antifade Mountant with DAPI (Invitrogen, P36935), coverslips were scanned on a SP8 confocal microscope. The fluorescent gray value of p-α-syn and MAP2 signal were measured by ImageJ.

### Cryo-EM sample preparation and data collection

The α-syn-PMCA derived fibrils at the concentration of 5 μM (4 μl) were added to the glow-discharged 300 meshed holey carbon copper Quantifoil grids (R2/1) twice. The grids were flash-frozen in liquid ethane after being blotted for 0.5 s by filter papers that were pretreated in Vitrobot Mark IV (FEI) with 95% humidity at 16 °C. The cryo-EM micrographs collection was accomplished by a Titan Krios transmission electron microscope (FEI) operated at 300 kV with a Gatan K3 camera at a magnification of 22,500x and in super-resolution mode. A range of -2.0 to -1.0 μm for defocus values was adopted. Each micrograph was dose-fractionated to 32 frames with the electron dose rate of ∼ 20 electrons per second per square pixel (∼ 20 e^-^/s/ pixel ^2^) and total dose of ∼ 55 e^-^/Å ^2^. Hence the total exposure time was set to 3.111 s (0.097 s/frame). All cryo-EM micrographs were collected automatically by Serial EM software.^45^

### Image pre-processing, helical reconstruction, and model building

MotionCorr2 was conducted for movie frames to correct beam induced motion and dose-weighted and micrographs with 32 frames were aligned, summed and further binned with a pixel size of 1.06 Å.^46^ Contrast transfer function were estimated using CTFFIND4.1.8.^47^ All fibrils were manually picked using the “ manual picking” program in RELION 3.1.^48^ All further processes of helical reconstruction were performed by RELION 3.1.

#### (1) Pre-PD dataset

45,846 pre-PD derived fibrils were manually picked from 2,567 micrographs. Initially, the pre-PD derived segments were extracted using a box size of 1024-pixel and an inter-box distance of 109 Å and used for subsequent reference-free 2D classification with a decreasing in-plane angular sampling rate from 12° to 1° and a T=2 regularization parameter to estimate the fibril pitch and helical parameters. Type 1A fibrils and Type 2 fibrils derived from the pre-PD were initially separated at this initial 2D classification. Due to feature of without twist, Type 2 fibrils subset could not be performed the further helical reconstruction. Type 1A fibrils containing selected segments with 1024-pixel box size were then re-extracted using a box size of 320-pixel and an inter-box distance of 34 Å and particles comprising an entire helical crossover were selected for following 3D classification. An *de novo* 3D initial model was constructed from selected particles after 2D classification by relion_helix_ inimodel2d program.^48^ *De novo* 3D initial model that was low-pass-filtered to 60 Å was further used as reference map of subsequent 3D classification. Optimization of helical twist and helical rise was conducted once β-strands showed a separation along the helix axis. Psedo-21 symmetry was applied and several rounds of 3D classification with K=3 and K=1 were performed to obtain homogeneous segments and optimize twist parameters. Optimized parameters and selected segments were applied for high-resolution gold-standard refinement. The final map of Type 1A fibrils was convergence with the rise of 2.44 Å and the twist angle of 179.45°. Post-processing with a soft-edge mask and an estimated map sharpening B-factor of -93.41 Å^2^ provided a map with a resolution of 2.78 Å according to the gold-standard FSC = 0.143 criteria. Atomic model of Type 1A fibrils was built into the central region of the helical reconstruction 3D density map with α-syn fibril polymorph 2b (PDB-ID: 6SST) used as guide in *Coot*.^10,49^ A model comprising three β-sheet rungs was generated and refined in PHENIX using real_space_refine.^50^

#### (2) Post-PD dataset

35,178 post-PD derived fibrils were manually picked from 2,594 micrographs. Initially, the post-PD derived segments were extracted using a box size of 686-pixel and an inter-box distance of 73 Å and used for subsequent reference-free 2D classification. Type 1B fibrils and Type 3 fibrils derived from the pre-PD were initially separated at this initial 2D classification. Type 1B fibrils containing selected segments with 686-pixel box size were then re-extracted using a box size of 288-pixel and an inter-box distance of 31 Å, while Type 3 fibrils containing selected segments with 686-pixel box size were then re-extracted using a box size of 224-pixel and an inter-box distance of 24 Å. Particles after 2D classification comprising an entire helical crossover were selected for following 3D classification. The construction of *de novo* 3D initial model, the selection of 3D reference map and the optimization of twist parameters for both Type 1B and Type 3 fibrils were performed as the “ pre-PD dataset” described. Several rounds of 3D classification with K=3 and K=1 was performed to accomplish the purification of segments and the optimization of twist parameters. Optimized parameters and homogeneous segments were applied for high-resolution gold-standard refinement. Post-processing with a soft-edge solvent mask was conducted to sharpen the refined maps. The final overall resolution estimate was calculated according to the 0.143 Fourier Shell Correlation cutoff. The atomic models building and refinement for Type 1B and Type 3 fibrils were performed as the “ pre-PD dataset” described. The atomic model with PDB-ID 6SST of α-syn fibril polymorph 2b and 7NCK of Type 3 α-syn fibrils amplified from MSA Case 5 were used as initial reference for Type 1B and Type 3 fibrils, respectively, in *Coot*.^10,51^

### Statistical analysis

Statistical significance in biological assays was measured by one-way ANOVA followed by Tukey’s post-hoc test using Prism version 7.0 (GraphPad Software, La Jolla, CA, United States). Values of *P* < 0.05 were considered statistically significant.

## Data availability statement

Cryo-EM 3D density maps were deposited in the Electron Microscopy Databank (EMDB) with entry codes EMD-31702 for Type 1A fibrils derived from the pre-PD, EMD-31703 for Type 1B fibrils and EMD-31704 Type 3 fibrils derived from the post-PD. The corresponding structure models were deposited in the Worldwide Protein Data Bank (wwPDB) with entry codes: 7V47, 7V48, and 7V49 for Type 1A, Type 1B and Type 3 fibrils, respectively. Additional data that support findings of this study will be available from the corresponding author upon reasonable request.

## Acknowledgments

This work was supported by the Major State Basic Research Development Program (Grant No. 2019YFE0120600), the National Natural Science Foundation (NSF) of China (Grant No. 91853113 31872716, 82171421, 91949118 and 81771372), the Science and Technology Commission of Shanghai Municipality (STCSM) (Grant No. 18JC1420500, 20XD1425000, 2019SHZDZX02 and 21S31902200), the “ Eastern Scholar” project supported by the Shanghai Municipal Education Commission, the Shanghai Municipal Science and Technology Major Project (Grant No. 2018SHZDZX01), ZJ Lab, and Shanghai Center for Brain Science and Brain-Inspired Technology.

## Author Contributions

D.L., J. W. and C.L. designed the project. Y.F. prepared the patients-derived fibril samples, performed the biochemical and cellular assays. Y.S. performed the cryo-EM experiments, built and refined the structure model. W.X. and Q.C. assisted in cryo-EM data collection and processing as well as model building. Y.S. and Y.T. assisted in figures preparation. W.Y., Y.L., Y.T. contributed to the clinical assessments of PD patients. Y.S., F.L. and J. W. contributed the CSF samples collection. All the authors are involved in analyzing the data and contributing to manuscript discussion and editing. Y.F and D.L. wrote the manuscript.

## Conflicts of interest

The authors declare no competing interests.

## Notes

### Competing Interest Statement

The authors have declared no competing interest.

## References

1 Wirdefeldt, K., Adami, H. O., Cole, P., Trichopoulos, D. & Mandel, J. Epidemiology and etiology of Parkinson’s disease: a review of the evidence. European journal of epidemiology 26, S1–S58, doi:10.1007/s10654-011-9581-6 (2011).

2 Spillantini, M. G. et al. Alpha-synuclein in Lewy bodies. Nature 388, 839–840, doi:10.1038/42166 (1997).

3 Spillantini, M. G., Crowther, R. A., Jakes, R., Hasegawa, M. & Goedert, M. alpha-Synuclein in filamentous inclusions of Lewy bodies from Parkinson’s disease and dementia with lewy bodies. Proceedings of the National Academy of Sciences of the United States of America 95, 6469–6473, doi:10.1073/pnas.95.11.6469 (1998).

4 Halliday, G. M., Holton, J. L., Revesz, T. & Dickson, D. W. Neuropathology underlying clinical variability in patients with synucleinopathies. Acta neuropathologica 122, 187–204, doi:10.1007/s00401-011-0852-9 (2011).

5 Bousset, L. et al. Structural and functional characterization of two alpha-synuclein strains. Nature communications 4, 2575, doi:10.1038/ncomms3575 (2013).

6 Lau, A. et al. α-Synuclein strains target distinct brain regions and cell types. Nature neuroscience 23, 21–31, doi:10.1038/s41593-019-0541-x (2020).

7 Peng, C. et al. Cellular milieu imparts distinct pathological α-synuclein strains in α-synucleinopathies. Nature 557, 558–563, doi:10.1038/s41586-018-0104-4 (2018).

8 Li, Y. et al. Amyloid fibril structure of α-synuclein determined by cryo-electron microscopy. Cell research 28, 897–903, doi:10.1038/s41422-018-0075-x (2018).

9 Guerrero-Ferreira, R. et al. Cryo-EM structure of alpha-synuclein fibrils. eLife 7, doi:10.7554/eLife.36402 (2018).

10 Guerrero-Ferreira, R. et al. Two new polymorphic structures of human full-length alpha-synuclein fibrils solved by cryo-electron microscopy. eLife 8, doi:10.7554/eLife.48907 (2019).

11 Li, B. et al. Cryo-EM of full-length α-synuclein reveals fibril polymorphs with a common structural kernel. Nature communications 9, 3609, doi:10.1038/s41467-018-05971-2 (2018).

12 Schweighauser, M. et al. Structures of α-synuclein filaments from multiple system atrophy. Nature 585, 464–469, doi:10.1038/s41586-020-2317-6 (2020).

13 Burger, D., Fenyi, A., Bousset, L., Stahlberg, H. & Melki, R. Cryo-EM structure of alpha-synuclein fibrils amplified by PMCA from PD and MSA patient brains. bioRxiv, 2021.2007.2008.451588, doi:10.1101/2021.07.08.451588 %J bioRxiv (2021).

14 Postuma, R. B. et al. MDS clinical diagnostic criteria for Parkinson’s disease. Movement disorders : official journal of the Movement Disorder Society 30, 1591–1601, doi:10.1002/mds.26424 (2015).

15 Tolosa, E., Garrido, A., Scholz, S. W. & Poewe, W. Challenges in the diagnosis of Parkinson’s disease. The Lancet. Neurology 20, 385–397, doi:10.1016/s1474-4422(21)00030-2 (2021).

16 Li, J. Y. et al. Lewy bodies in grafted neurons in subjects with Parkinson’s disease suggest host-to-graft disease propagation. Nature medicine 14, 501–503, doi:10.1038/nm1746 (2008).

17 Masuda-Suzukake, M. et al. Prion-like spreading of pathological α-synuclein in brain. Brain : a journal of neurology 136, 1128–1138, doi:10.1093/brain/awt037 (2013).

18 Prusiner, S. B. et al. Evidence for α-synuclein prions causing multiple system atrophy in humans with parkinsonism. Proceedings of the National Academy of Sciences of the United States of America 112, E5308–5317, doi:10.1073/pnas.1514475112 (2015).

19 Olanow, C. W., Stern, M. B. & Sethi, K. The scientific and clinical basis for the treatment of Parkinson disease (2009). Neurology 72, S1–136, doi:10.1212/WNL.0b013e3181a1d44c (2009).

20 Braak, H. et al. Staging of brain pathology related to sporadic Parkinson’s disease. Neurobiology of aging 24, 197–211, doi:10.1016/s0197-4580(02)00065-9 (2003).

21 Dunning, C. J., Reyes, J. F., Steiner, J. A. & Brundin, P. Can Parkinson’s disease pathology be propagated from one neuron to another? Progress in neurobiology 97, 205–219, doi:10.1016/j.pneurobio.2011.11.003 (2012).

22 Shahnawaz, M. et al. Development of a Biochemical Diagnosis of Parkinson Disease by Detection of α-Synuclein Misfolded Aggregates in Cerebrospinal Fluid. JAMA neurology 74, 163–172, doi:10.1001/jamaneurol.2016.4547 (2017).

23 He, S. & Scheres, S. H. W. Helical reconstruction in RELION. Journal of structural biology 198, 163–176, doi:10.1016/j.jsb.2017.02.003 (2017).

24 Li, D. & Liu, C. Hierarchical chemical determination of amyloid polymorphs in neurodegenerative disease. Nature chemical biology 17, 237–245, doi:10.1038/s41589-020-00708-z (2021).

25 Fitzpatrick, A. W. P. et al. Cryo-EM structures of tau filaments from Alzheimer’s disease. Nature 547, 185–190, doi:10.1038/nature23002 (2017).

26 Falcon, B. et al. Novel tau filament fold in chronic traumatic encephalopathy encloses hydrophobic molecules. Nature 568, 420–423, doi:10.1038/s41586-019-1026-5 (2019).

27 Zhang, W. et al. Novel tau filament fold in corticobasal degeneration. Nature 580, 283–287, doi:10.1038/s41586-020-2043-0 (2020).

28 Frieg, B. et al. α-Synuclein polymorphism determines oligodendroglial dysfunction. BioRxiv, 2021.2007.2009.451731, doi:10.1101/2021.07.09.451731 %J bioRxiv (2021).

29 Zhao, K. et al. Parkinson’s disease-related phosphorylation at Tyr39 rearranges α-synuclein amyloid fibril structure revealed by cryo-EM. Proceedings of the National Academy of Sciences of the United States of America 117, 20305–20315, doi:10.1073/pnas.1922741117 (2020).

30 Falcon, B. et al. Tau filaments from multiple cases of sporadic and inherited Alzheimer’s disease adopt a common fold. Acta neuropathologica 136, 699–708, doi:10.1007/s00401-018-1914-z (2018).

31 Shi, Y. et al. Structure-based Classification of Tauopathies. BioRxiv, 2021.2005.2028.446130, doi:10.1101/2021.05.28.446130 %J bioRxiv (2021).

32 Xiang, X. et al. Role of molecular polymorphism in defining tau filament structures in neurodegenerative diseases. BioRxiv, 2021.2005.2024.445353, doi:10.1101/2021.05.24.445353 %J bioRxiv (2021).

33 Falcon, B. et al. Structures of filaments from Pick’s disease reveal a novel tau protein fold. Nature 561, 137–140, doi:10.1038/s41586-018-0454-y (2018).

34 Chaudhuri, K. R. et al. International multicenter pilot study of the first comprehensive self-completed nonmotor symptoms questionnaire for Parkinson’s disease: The NMSQuest study. Movement Disorders 21, 916–923, doi:10.1002/mds.20844 (2006).

35 Ertan, F. S., Ertan, T., Kiziltan, G. & Uyguçgil, H. Reliability and validity of the Geriatric Depression Scale in depression in Parkinson’s disease. Journal of neurology, neurosurgery, and psychiatry 76, 1445–1447 (2005).

36 Richter, P., Werner, J., Heerlein, A., Kraus, A. & Sauer, H. On the validity of the Beck Depression Inventory. A review. Psychopathology 31, 160–168, doi:10.1159/000066239 (1998).

37 Wang, Y. et al. Validation of the rapid eye movement sleep behavior disorder screening questionnaire in China. J Clin Neurosci 22, 1420–1424, doi:10.1016/j.jocn.2015.03.008 (2015).

38 Chen, N.-H. et al. Validation of a Chinese version of the Epworth sleepiness scale. Qual Life Res 11, 817–821 (2002).

39 Katzman, R. et al. A Chinese version of the Mini-Mental State Examination; impact of illiteracy in a Shanghai dementia survey. J Clin Epidemiol 41, 971–978 (1988).

40 Tsang, K.-L. et al. Translation and validation of the standard Chinese version of PDQ-39: a quality-of-life measure for patients with Parkinson’s disease. Movement disorders : official journal of the Movement Disorder Society 17, 1036–1040 (2002).

41 Zhao, H. et al. AlphaLISA detection of alpha-synuclein in the cerebrospinal fluid and its potential application in Parkinson’s disease diagnosis. Protein & cell 8, 696–700, doi:10.1007/s13238-017-0424-4 (2017).

42 Johnson, M., Coulton, A. T., Geeves, M. A. & Mulvihill, D. P. Targeted amino-terminal acetylation of recombinant proteins in E. coli. PloS one 5, e15801, doi:10.1371/journal.pone.0015801 (2010).

43 Kam, T. I. et al. Poly(ADP-ribose) drives pathologic α-synuclein neurodegeneration in Parkinson’s disease. Science (New York, N.Y.) 362, doi:10.1126/science.aat8407 (2018).

44 Shahnawaz, M. et al. Discriminating α-synuclein strains in Parkinson’s disease and multiple system atrophy. Nature 578, 273–277, doi:10.1038/s41586-020-1984-7 (2020).

45 Mastronarde, D. N. Automated electron microscope tomography using robust prediction of specimen movements. Journal of structural biology 152, 36–51, doi:10.1016/j.jsb.2005.07.007 (2005).

46 Zheng, S. Q. et al. MotionCor2: anisotropic correction of beam-induced motion for improved cryo-electron microscopy. Nature methods 14, 331–332, doi:10.1038/nmeth.4193 (2017).

47 Rohou, A. & Grigorieff, N. CTFFIND4: Fast and accurate defocus estimation from electron micrographs. Journal of structural biology 192, 216–221, doi:10.1016/j.jsb.2015.08.008 (2015).

48 Scheres, S.H.W. Amyloid structure determination in RELION-3.1. Acta crystallographica. Section D, Structural biology 76, 94–101, doi:10.1107/s2059798319016577 (2020).

49 Emsley, P., Lohkamp, B., Scott, W. G. & Cowtan, K. Features and development of Coot. Acta crystallographica. Section D, Biological crystallography 66, 486–501, doi:10.1107/s0907444910007493 (2010).

50 Adams, P. D. et al. PHENIX: a comprehensive Python-based system for macromolecular structure solution. Acta crystallographica. Section D, Biological crystallography 66, 213–221, doi:10.1107/s0907444909052925 (2010).

51 Lövestam, S. et al. Seeded assembly in vitro does not replicate the structures of α-synuclein filaments from multiple system atrophy. FEBS open bio 11, 999–1013, doi:10.1002/2211-5463.13110 (2021).

