## Supplementary Figure 1/2/3/4/5 for "Different structures and pathologies of α-synuclein fibrils derived from preclinical and postmortem patients of Parkinson’s disease"

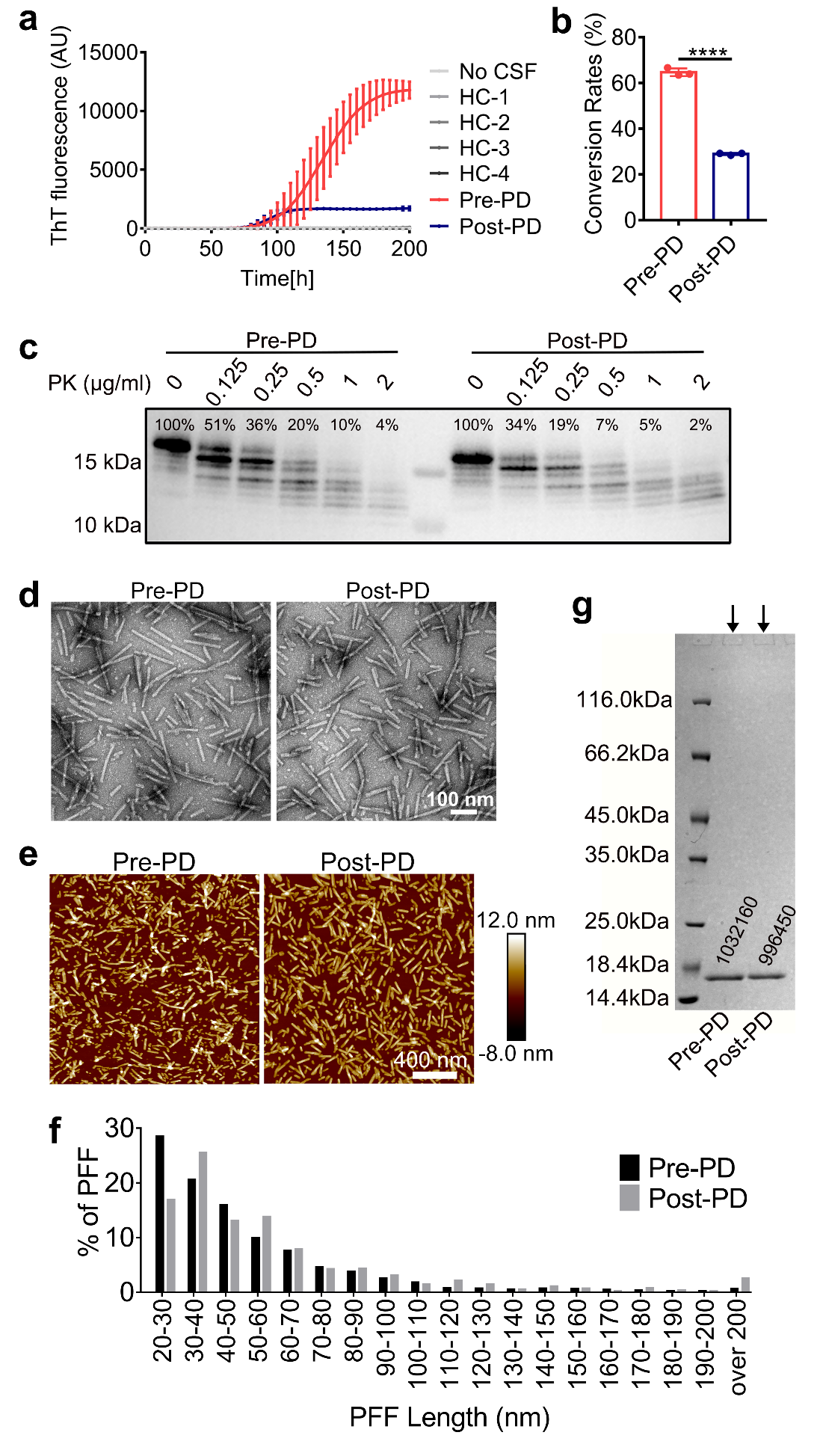

**Supplementary Figure 1. Amplification and characterization of α-syn fibrils. a.** ThT kinetic assay forα-syn aggregation seeded with CSF from healthy controls (HCs), a pre-PD patient and a late-stage post-PD patient. Same result as shown in Fig. 1a with y-axis presenting the intensity of ThT fluorescence. Data shown are mean ± SD, n=3 individual experimental samples. **b**. The conversion rates of α-syn monomer to fibrils in pre-PD and post-PD samples in (**a**). Data shown are mean ± SD, n=3. **c**. Western blot of pre-PD and post-PD PFFs digested by PK. Fibrils amplified from pre-PD and post-PD CSF were sonicated to prepare PFFs. The PFFs were incubated with the indicated concentration of PK at 37 °C for 30 min. The percentage of full-length α-syn in each lane was calculated based on the band intensities measured by Image J. **d.** NS-TEM images of pre-PD and post-PD α-syn PFFs. **e.** AFM images of pre-PD and post-PD α-syn PFFs. **f**. Lengths of α-syn PFFs measured by AFM. 2,341 pre-PD PFFs and 2,280 post-PD PFFs were measured, respectively. **g**. SDS-PAGE gel of the pre-PD and post-PD α-syn PFFs used to treat primary neurons. The loading holes of the gel are indicated by arrows and the band intensities were analyzed by Image J.

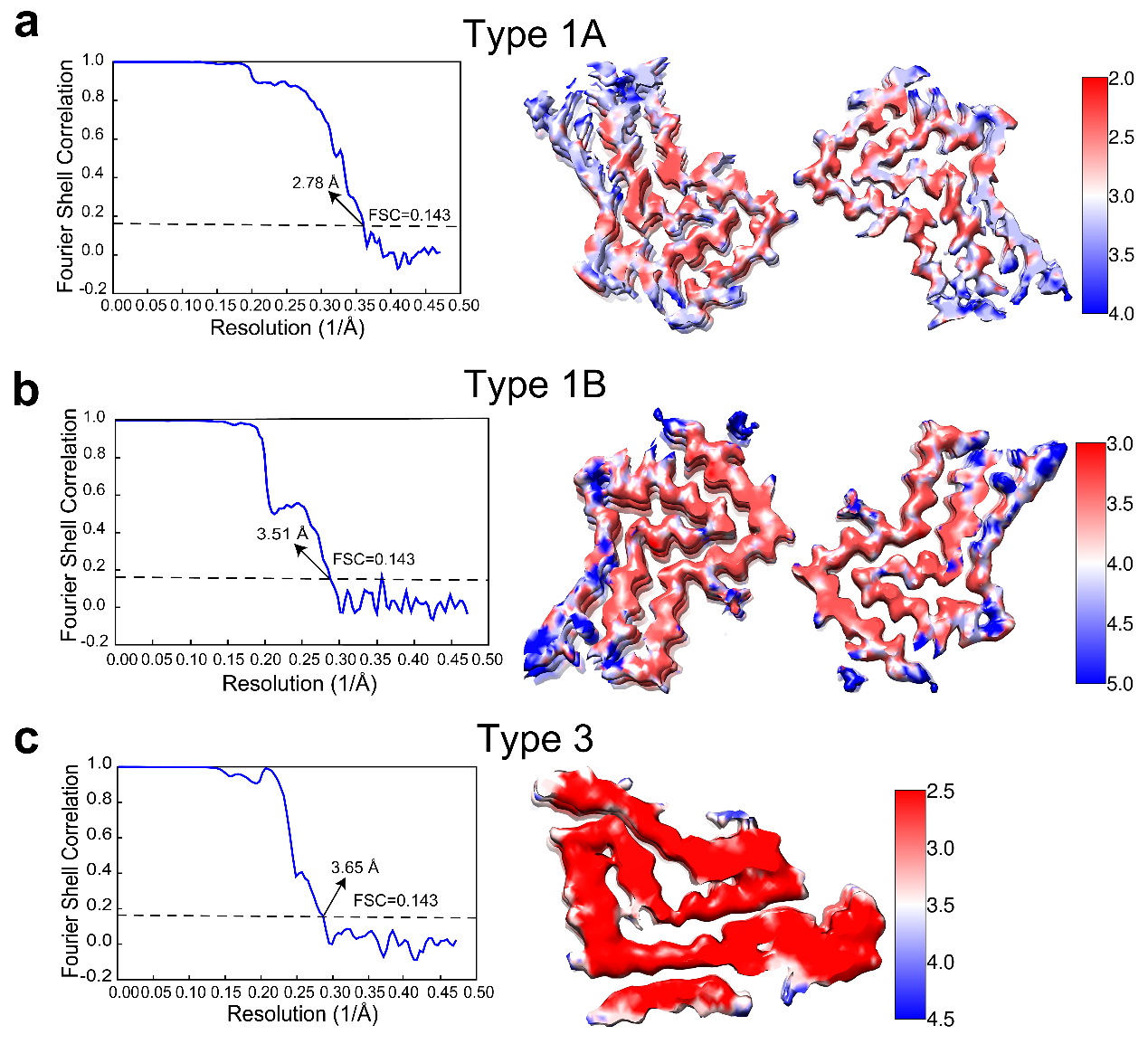

**Supplementary Figure 2. Resolution estimation of the cryo-EM structures of the pre-PD and post-PD derived fibrils.** Gold standard Fourier shell correction curves (left) and local resolution estimations of the reconstruction (right) for the Type 1A fibril **(a),** Type 1B fibril **(b),** and Type 3 fibril **(c)**. The overall resolution of the Type 1A, Type 1B, Type 3 fibrils is 2.78 Å, 3.51 Å, and 3.65 Å, respectively.

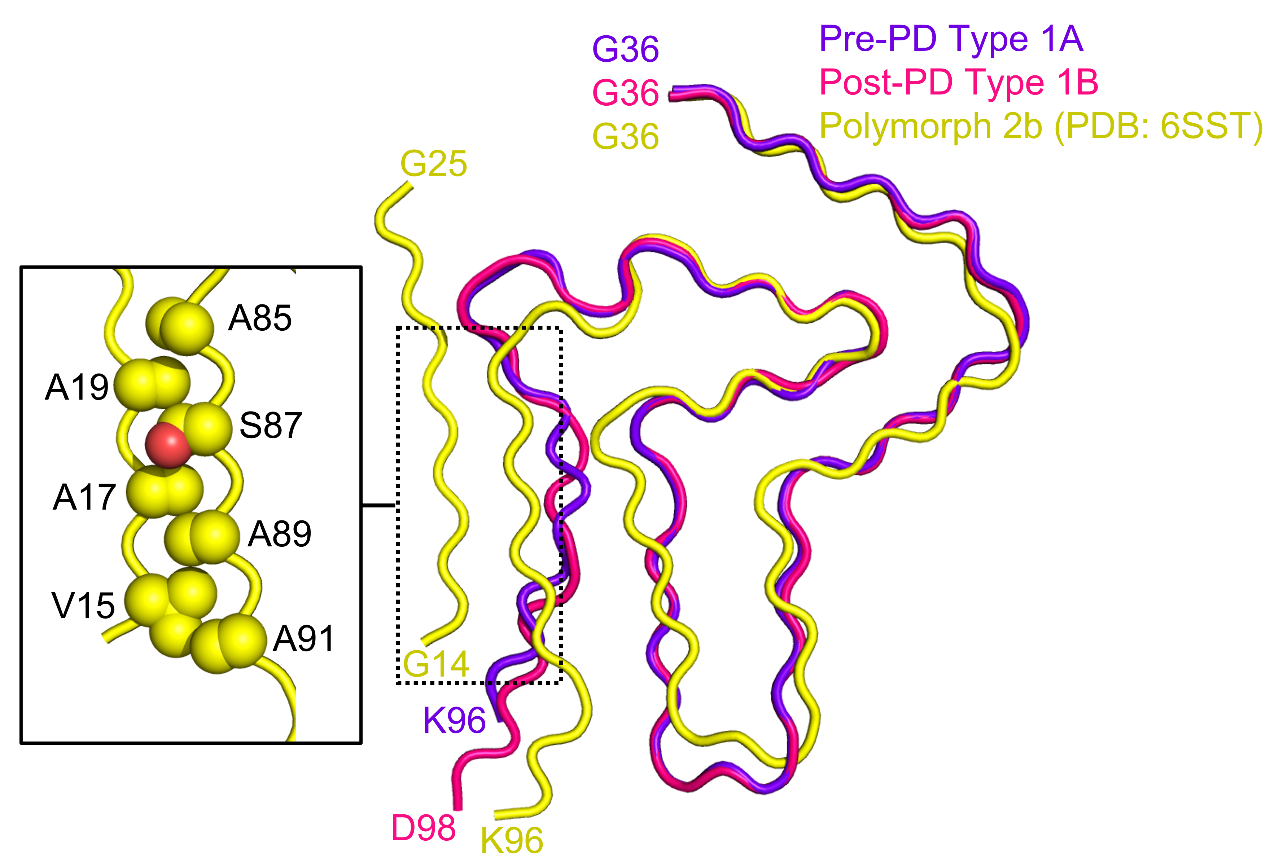

**Supplementary Figure 3. Structural comparisons of the Type 1A and Type 1B α-syn fibrils amplified from PD CSF, and the polymorph 2b α-syn fibril.** Overlay of α-syn monomer structures in the Type 1A, Type 1B and polymorph 2b (PDB ID: 6SST) fibrils. The steric-zipper-like interaction between residues 15-19 and 85-91 in polymorph 2b is shown in the zoom-in box. Side chains are shown as spheres.

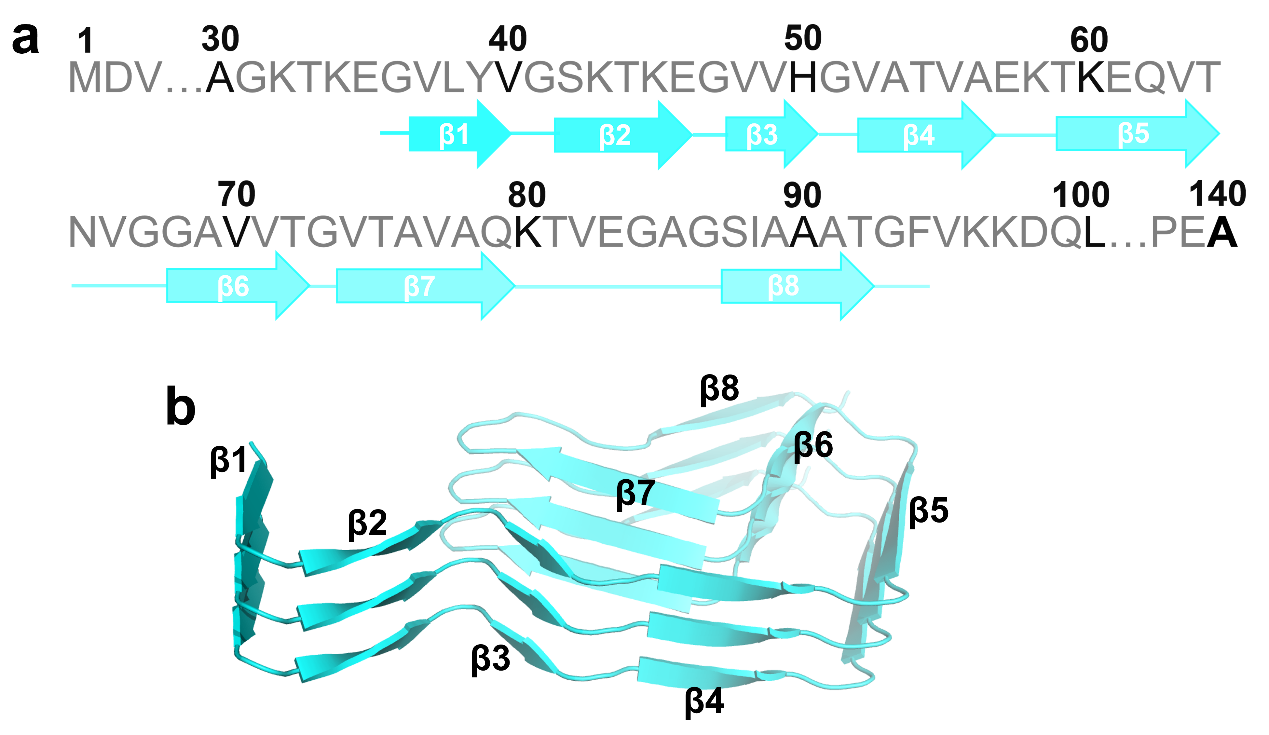

**Supplementary Figure 4. Cartoon views of the secondary structural elements in the Type 3 fibril. a.** Primary sequence of human wild type α-syn with the β strands of Type 3 fibril indicated by arrows. **b.** Cartoon views of the secondary structural elements in three successive rungs of Type 3 fibril with β strands labeled.

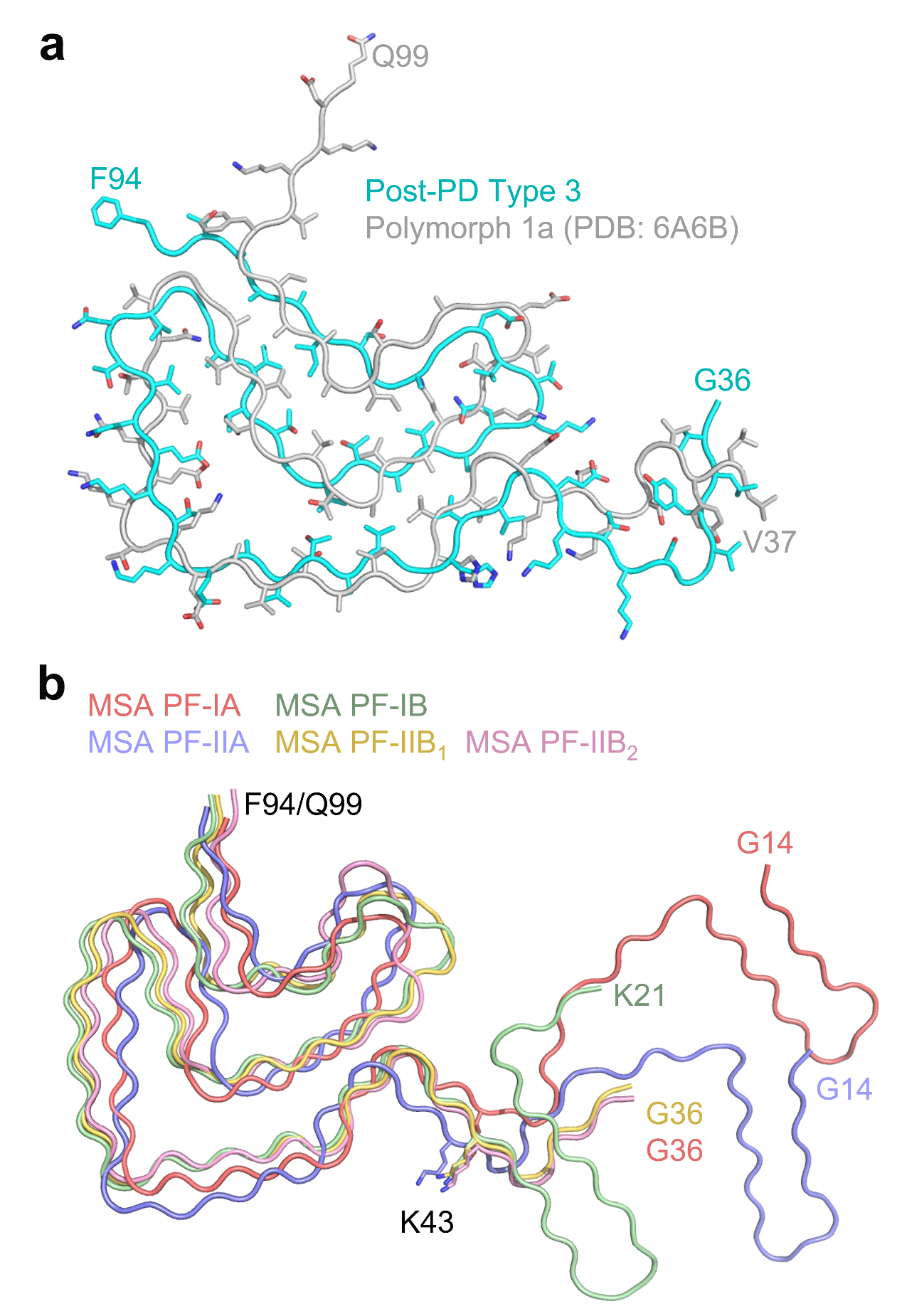

**Supplementary Figure 5. Structural comparisons of polymorphic α-syn fibrils. a.** Overlay of the α-syn structures in the post-PD Type 3 fibril and the polymorph 1a fibril. The RMSD between the two structures is 4.85 Å over 58 C-α atoms. Side chains are shown as sticks. **b.** Overlay of α-syn polymorphic structures in MSA-purified fibrils. Residues K43-F94/Q99 represent a conserved structural motif. Side chain of residue K43 is shown as sticks.
